# Stress granule-inducing eukaryotic translation initiation factor 4A inhibitors block influenza A virus replication

**DOI:** 10.1101/194589

**Authors:** Patrick D. Slaine, Mariel Kleer, Nathan Smith, Denys A. Khaperskyy, Craig McCormick

## Abstract

Eukaryotic translation initiation factor 4A (eIF4A) is a helicase that facilitates assembly of the translation preinitiation complex by unwinding structured mRNA 5’ untranslated regions. Pateamine A (PatA) and silvestrol are natural products that disrupt eIF4A function and arrest translation, thereby triggering the formation of cytoplasmic aggregates of stalled preinitiation complexes known as stress granules (SGs). Here we examined the effects of eIF4A inhibition by PatA and silvestrol on influenza A virus (IAV) protein synthesis and replication in cell culture. Treatment of infected cells with either PatA or silvestrol at early times post-infection results in SG formation, arrest of viral protein synthesis and failure to replicate the viral genome. PatA, which irreversibly binds to eIF4A, sustained long-term blockade of IAV replication following drug withdrawal, and inhibited IAV replication at concentrations that had minimal cytotoxicity. By contrast, the antiviral effects of silvestrol were fully reversible; drug withdrawal caused rapid SG dissolution and resumption of viral protein synthesis. IAV inhibition by silvestrol was invariably associated with cytotoxicity. PatA blocked replication of genetically divergent IAV strains, suggesting common dependence on host eIF4A activity. This study demonstrates the feasibility of targeting core host protein synthesis machinery to prevent viral replication.

**IMPORTANCE:** Influenza A virus (IAV) relies on cellular protein synthesis to decode viral messenger RNAs. Pateamine A and silvestrol are natural products that inactivate an essential protein synthesis protein known as eIF4A. Here we show that IAV is sensitive to these eIF4A inhibitor drugs. Treatment of infected cells with pateamine A or silvestrol prevented synthesis of viral proteins, viral genome replication and release of infectious virions. The irreversible eIF4A inhibitor pateamine A sustained long-term blockade of viral replication, whereas viral protein synthesis quickly resumed after silvestrol was removed from infected cells. Prolonged incubation of either infected or uninfected cells with these drugs induced the programmed cell death cascade called apoptosis. Our findings suggest that core components of the host protein synthesis machinery are viable targets for antiviral drug discovery. The most promising drug candidates should selectively block protein synthesis in infected cells without perturbing bystander uninfected cells.

## INTRODUCTION

Influenza A viruses (IAV) are enveloped viruses with segmented, negative-sense single-stranded RNA genomes (vRNAs) that primarily infect epithelial cells in in a diverse range of hosts. The viruses bind to cell surface sialic acids and are internalized by endocytosis. The virus gains access to the cytoplasm following the fusion of viral and endosome membranes. Viral genome segments are then imported into the cell nucleus, where viral mRNA synthesis is initiated by the viral RNA-dependent RNA polymerase (RdRp) complex, which cleaves nascent host RNA polymerase II (pol II)-transcribed RNAs at sites 10-14 nucleotides downstream of 5’ 7-methyl-guanosine (m7G) caps (1). These short capped RNAs are used to prime viral mRNA synthesis. Template-directed viral mRNA synthesis is terminated by reiterative decoding of short uridine-rich sequences, which generates 3’-poly-adenylate (polyA) tails. Thus, by coupling a trimeric RdRp complex to each incoming genome segment, the virus ensures faithful generation of mRNAs with 5’-caps and polyA tails that largely resemble host mRNAs.

Efficient translation of IAV mRNAs depends on virus-mediated suppression of host gene expression, a process known as host shutoff. Host shutoff gives viral mRNAs priority access to the host protein synthesis machinery. IAV host shutoff mechanisms identified to date encompass different stages of host mRNA biogenesis. The viral PA-X endoribonuclease selectively targets host pol II transcripts while sparing viral mRNAs, as well as pol I and pol III transcripts (2). In addition, the viral nonstructural-1 (NS1) protein binds and inhibits cleavage and polyadenylation specificity factor 30 kDa subunit (CPSF30), an essential component of the host 3’ end processing machinery for cellular pre-mRNAs that is dispensable for viral mRNA processing (3). NS1 also enhances the association between newly-synthesized viral mRNAs and cellular NXF1/Tap mRNA export factors (4), while simultaneously inhibiting nucleoporin 98 (Nup98)-mediated export of host mRNAs (5). In addition to host shutoff, there is evidence that efficient translation of viral mRNAs depends on short, conserved cis-acting sequences in viral 5’ untranslated regions (UTRs), downstream of the host-derived leader sequences (6). It is not known precisely how these 5’-UTR sequences assist translation, but recent work has shown that when bound to viral mRNAs, the NS1 protein plays an important role in ribosome recruitment (7). By contrast, recent studies employing RNA-sequencing, ribosomal foot-printing and single-molecule fluorescence in-situ hybridization have suggested that host shutoff is mainly achieved by reduction in cellular mRNA levels, and that IAV mRNAs are not preferentially translated (8). It is clear that viral proteins enable efficient translation of IAV mRNAs, but the precise composition of translation pre-initiation complexes assembled on these mRNAs, and the contribution of diverse host translation initiation factors remains incompletely characterized.

Dependence on host protein synthesis machinery makes viral mRNAs sensitive to various stress-induced translation repression mechanisms. A particularly critical checkpoint in translation initiation is the assembly of the ternary complex, comprised of eIF2, GTP and tRNAi^met^, which allows for incorporation of the tRNAi^met^ into the 40S ribosomal subunit. This step is negatively regulated by serine phosphorylation of eIF2□by one of four kinases: heme-regulated translation inhibitor (HRI), general control non-derepressible-2 (GCN2), protein kinase R (PKR) and PKR-like endoplasmic reticulum kinase (PERK) (9). eIF2□phosphorylation causes accumulation of stalled 48S preinitiation complexes and associated mRNA transcripts, which are bound by aggregation-prone RNA-binding proteins, including Ras-GAP SH3 domain-binding protein (G3BP), T-cell intracellular antigen-1 (TIA-1), and TIA-1 related protein (TIAR). These complexes nucleate cytoplasmic stress granules (SGs), sites where stalled mRNP complexes are triaged, until stress is resolved and protein synthesis can resume.

IAV mRNAs are efficiently translated throughout infection and SGs never form (10). IAV prevents translation arrest, at least in part, through the action of NS1, a double-stranded RNA (dsRNA)-binding protein that prevents PKR-mediated eIF2α phosphorylation. We have shown previously that IAV encodes two additional proteins, PA-X and NP, that block SG formation in an eIF2□-independent manner (11). Precise details of NP and PA-X mechanism of action remain obscure, but SG-inhibition by PA-X is tightly linked to its host shutoff function, as it requires endonuclease activity. This mechanism is reminiscent of the herpes simplex virus-type 2 (HSV-2) virion host shutoff (vhs) protein, which also requires endonuclease activity to prevent SG formation (12). Taken together, these findings suggest that IAV dedicates a significant portion of its small genome to encode proteins with SG-antagonizing activity.

SG suppression is maximal at late times post-infection, coinciding with the accumulation of NS1, NP and PA-X proteins. We have shown previously that at early times post-infection, treatment of infected cells with sodium arsenite, which causes HRI activation and eIF2α phosphorylation, triggered SG formation and stalled viral replication (11). Thus, like host mRNAs, at early times post-infection, IAV mRNAs are exquisitely sensitive to eIF2α phosphorylation and ternary complex depletion. We have also shown that IAV mRNAs are sensitive to pateamine A (PatA) (11), a natural product that selectively inhibits the DEAD-box RNA helicase eIF4A which assembles with eIF4E and eIF4G, to form the eIF4F complex (13, 14). By disrupting eIF4A, PatA prevents scanning by the 43S pre-initiation complex, causing arrest of translation initiation, and inducing SG formation (15). Specifically, treatment with PatA caused SG formation at early times post-infection and diminished accumulation of viral proteins.

In addition to PatA, eIF4A can be inhibited by a variety of natural compounds currently being investigated for anti-cancer properties, including hippuristanol (16), 15-deoxy-delta 12,14-prostaglandin J2 (15d-PGJ2) (17), silvestrol (18) and synthetic derivatives. Accumulating evidence indicates that many viruses are highly dependent on eIF4A activity for protein synthesis. For example, PatA has been shown to limit human cytomegalovirus infection (19), whereas silvestrol has been shown to inhibit Ebola virus replication (20).

In this study, we methodically documented the antiviral properties of PatA and silvestrol in three workhorse models of IAV infection; A549 human lung adenocarcinoma cells, Madin-Darby canine kidney (MDCK) cells, and African green monkey kidney epithelial cells (Vero). In all cases, dosages of PatA and silvestrol sufficient to elicit SGs concurrently blocked viral protein accumulation and genome replication. PatA, known to bind irreversibly to eIF4A, could sustain long-term arrest of viral protein synthesis following drug withdrawal, whereas the effects of silvestrol were reversible. Finally, PatA could inhibit genetically-divergent strains, suggesting common dependence on eIF4A.

## MATERIALS AND METHODS

### Reagents and cells

Unless specifically indicated, all chemicals were purchased from Sigma. Pateamine A was a kind gift from Dr. Jerry Pelletier (McGill University, Montreal, QC, Canada), silvestrol was obtained from MedChem Express (Princeton, NJ); 1 mM stocks of silvestrol and pateamine A were prepared in DMSO and stored at -80C. A549, Vero and Madin-Darby Canine Kidney (MDCK) cells were maintained in Dulbecco’s modified Eagle’s medium (DMEM; HyClone) supplemented with 10% fetal bovine serum (FBS, Life Technologies) and 100 U/ml penicillin + 100 μg/ml streptomycin + 20 μM L-glutamine (Pen/Strep/Gln; Wisent) at 37°C in 5% CO2 atmosphere. All cell lines were purchased from the American Type Culture Collection (ATCC).

### Viruses, infections and treatments

Influenza virus strains used in this study include A/PuertoRico/8/34/(H1N1) virus (PR8) and A/Udorn/1972(H3N2) virus (Udorn). Unless specified otherwise, infections were conducted at a multiplicity of infection (MOI) of 1. After inoculation, cells were washed with 1x PBS and incubated with infection medium containing 0.5% bovine serum albumin (BSA) in DMEM and incubated at 37°C in 5% CO2 atmosphere. IAV virions were enumerated by plaque assay using 1.2% Avicel RC 591 (FMC Corporation) overlay on confluent MDCK cells as described in (21). Where indicated, mock and virus-infected cells were treated with 0.156-40 nM pateamine A or 2.5-640 nM silvestrol.

### Cell viability measurement

Cells were seeded 24 h before drug treatment at a density of 10,000 cells/well in 96-well plates. Drugs were serially-diluted in media to the indicated concentrations and added to cells. After 24-h treatment, AlamarBlue assay (Thermo Scientific) was conducted according to manufacturer’s protocol using Tecan Infinite M200 PRO microplate reader (excitation = 560 nm, emission = 590 nm). Values were normalized to vehicle control.

### Immunostaining

For immunofluorescence microscopy cells grown on glass coverslips were fixed and stained as described previously (11) using mouse monoclonal antibody to G3BP (clone 23, BD Transduction Labs), goat polyclonal antibody to influenza virus (ab20841, Abcam), and rabbit monoclonal antibody to TIAR (clone D32D3, Cell Signaling) at manufacturer-recommended dilutions. Donkey Alexa Fluor-conjugated secondary antibodies (Molecular Probes) were used at 1:1000 dilution in combination with 5 ng/ml Hoechst dye. Images were captured using Zeiss Axioplan II microscope or Zeiss LSM 510 laser scanning microscope. For western blot analysis, whole cell lysates were resolved on TGX Stain-Free Precast Gels (BioRad) and analyzed using goat polyclonal antibody to influenza virus described above (recognizes HA1, NP, and M1 proteins), mouse monoclonal antibodies to influenza NS1 (clone 13D8, reference (22)), puromycin (MABE343, Millipore Sigma), and rabbit antibodies to phospho-Ser-51-eIF2α (D9G8, Cell Signaling), PARP (9542S, Cell Signaling), and β-actin (13E5, HRP-conjugated, Cell Signaling).

### Real time quantitative PCR

Total RNA was isolated using the RNeasy Plus Mini kit (Qiagen) according to manufacturer’s protocol. Viral genomic and messenger RNAs were reverse-transcribed as described in (2) using Maxima H Minus Reverse Transcriptase (Thermo Scientific) in separate reactions containing the gene-specific primer for 18S rRNA (5’-AGG GCC TCA CTA AAC CAT CC-3’) and either the influenza A virus-specific universal primer Uni12 (5’-AGC AAA AGC AGG-3’, for vRNA) or the oligo(dT)_18_ primer (for mRNA). Quantitative PCR analysis was performed using GoTaq PCR master mix (Promega). Relative initial template quantities were determined using the Ct method. Primer sequences and the PCR thermal profile setup are available upon request.

## RESULTS

### Pateamine A and silvestrol induce stress granules and inhibit viral protein accumulation in dose-dependent manner

We previously demonstrated that IAV was sensitive to pharmacologic induction of translation arrest and SG formation in the early stages of infection (11). To gain a fuller understanding of how eIF4A-targeting drugs can trigger SG formation and disrupt viral replication, IAV-infected A549, Vero and MDCK cells were treated at 1 hpi with increasing doses of PatA and silvestrol. In all three cell lines, PatA triggered SG formation and inhibited IAV protein accumulation at concentrations above 2.5 nM (Fig. 1A-D). By contrast, SG formation was triggered in response to 40 nM silvestrol in infected A549 cells, and 160 nM silvestrol in infected MDCK cells. Remarkably, Vero cells were highly resistant to translation arrest and SG induction by silvestrol. For both PatA and silvestrol, SG induction tightly correlated with inhibition of viral protein synthesis across all three cell lines.

**Figure 1.**
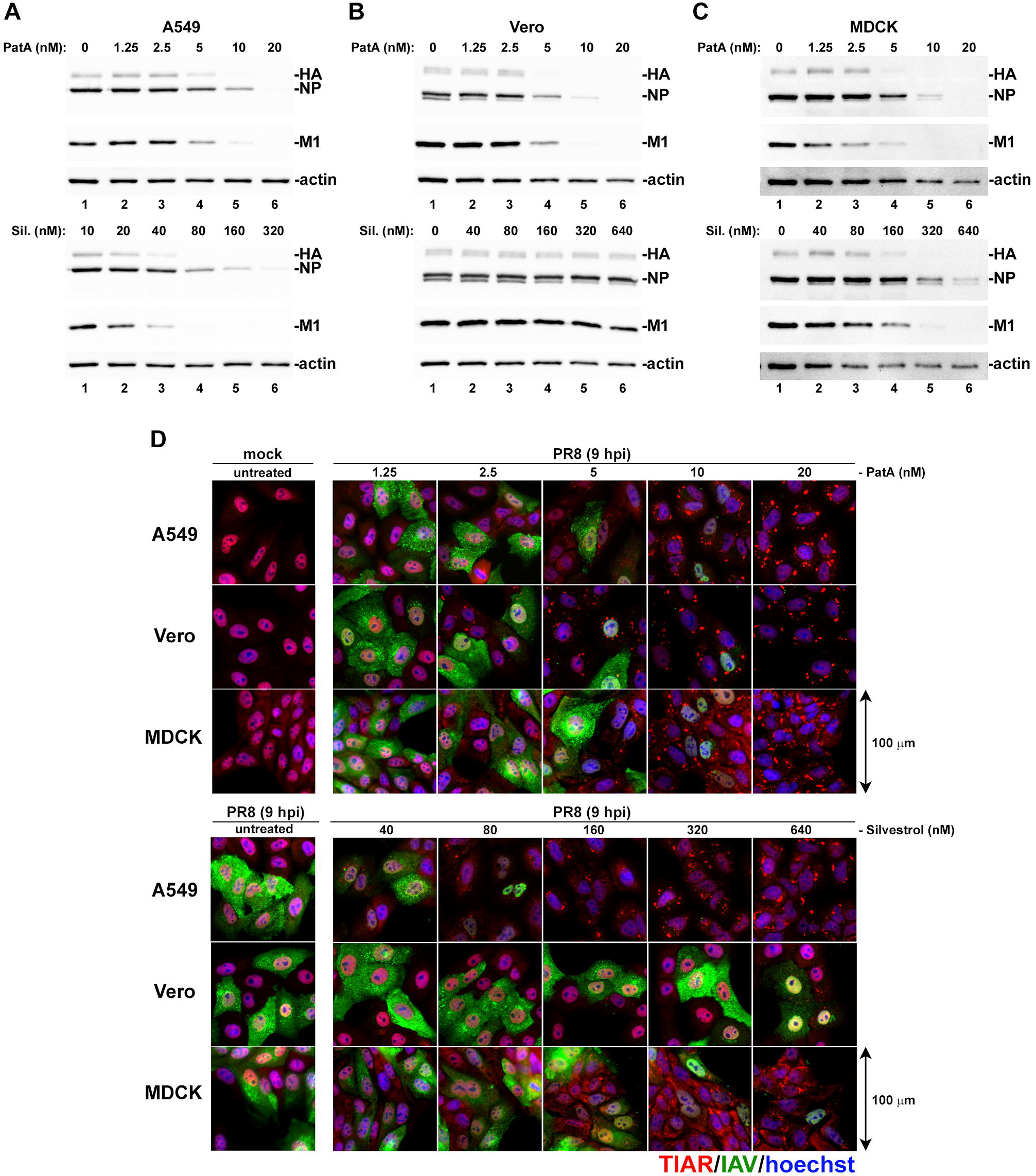
Concentration-dependent stress granule induction and inhibition of viral protein accumulation by pateamine A and silvestrol. A549, Vero, and MDCK cell lines infected with PR8 strain of IAV were treated with the indicated concentrations of pateamine A (PatA) or silvestrol (Sil.) at 1 hpi. (A-C) Viral protein accumulation was analysed in whole cell lysates collected at 24 hpi using western blot. (D) Stress granule induction and viral protein accumulation were visualized at 9 hpi by immunofluorescence microscopy using antibody to stress granule marker TIAR (red) and the polyclonal anti-influenza antibody (green). Nuclei were stained with Hoechst dye (blue). Representative images are shown for each cell line and treatment condition.

### Pateamine A is more potent than silvestrol at inhibiting viral replication in cultured cells

Previously, we demonstrated that 10 nM PatA blocked IAV replication in A549 cells through sustained total protein synthesis arrest and SG formation (11). To further test the effects of PatA and silvestrol on infectious virus release in the three most common cell culture models of IAV infection, we treated A549, Vero and MDCK cells with increasing concentrations of these drugs at 1 h post-infection with the PR8 strain of IAV. At 24 hpi, culture supernatants were harvested and the infectious virus titers were determined using plaque assays in MDCK cells (Fig. 2A, B). In agreement with results presented in Fig. 1, 5 nM PatA was sufficient to reduce virus replication 10-fold in all three cell lines while silvestrol had minimal effect on virus replication in Vero cells. In MDCK cells, silvestrol was much less effective at inhibiting infectious virus release;16-times higher drug concentration was required to match the inhibition of infectious virion release observed in A549 cells. Overall, these results reveal tight correlation between viral protein accumulation and the infectious virus production in these cell lines, and the unexpected remarkable resistance of Vero cells to silvestrol. Next, we tested the effects of PatA and silvestrol on cell viability after 24 h treatment using the AlamarBlue assay (Fig. 2C, D). Both drugs exhibited notable cytotoxicity at the concentrations required for maximal inhibition of virus replication. However, the cytotoxic effects varied significantly between the cell lines tested. 5 nM PatA had little effect on A549 cells, but it caused a sharp decrease in viability of MDCK cells. By contrast, among the three cell types, A549 cells were most sensitive to silvestrol. Taken altogether, these results suggest that PatA is more effective than silvestrol at inhibiting virus replication at sub-cytotoxic concentrations and exhibits a narrow therapeutic index in A549 cells.

**Figure 2.**
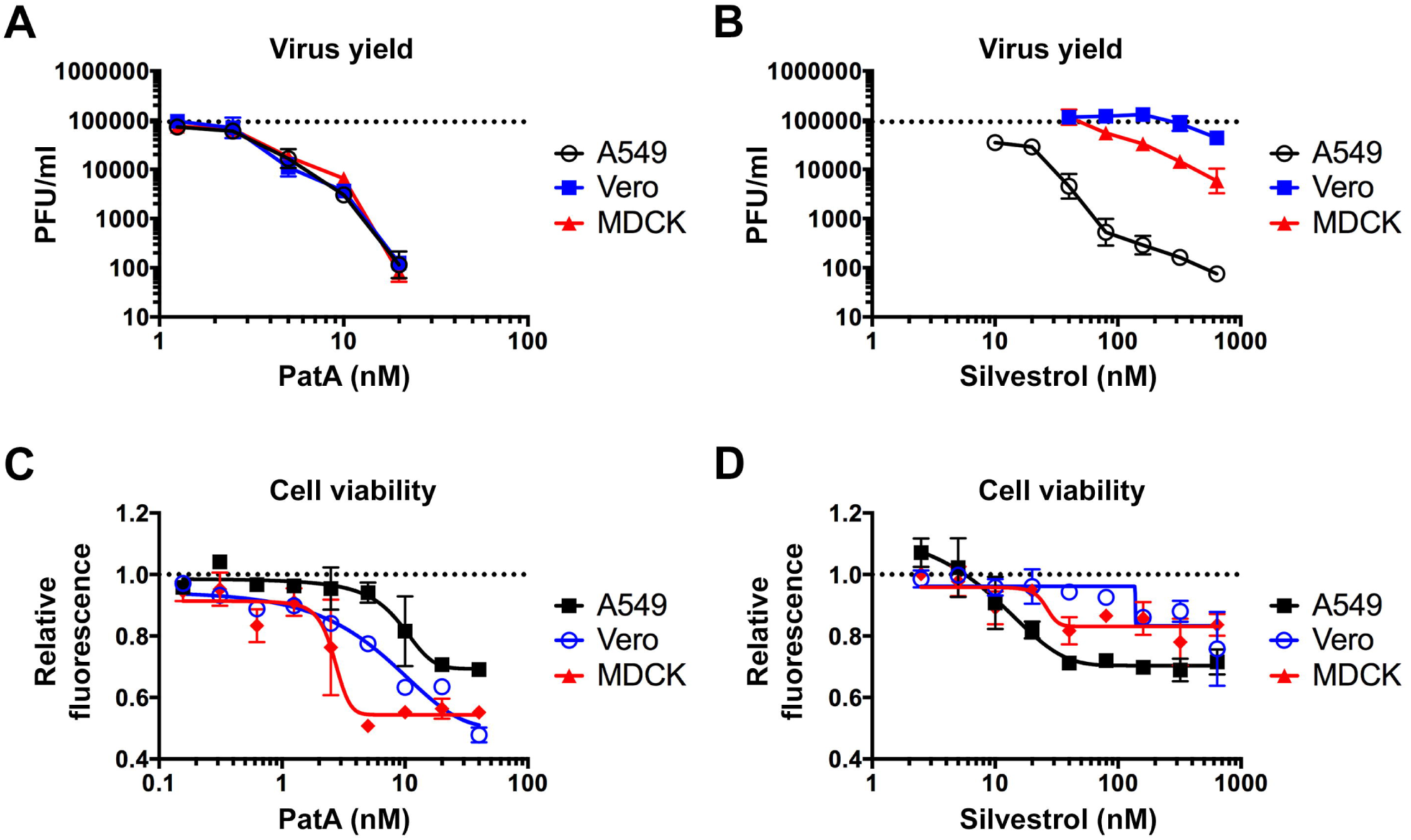
Antiviral and cytotoxic effects of pateamine A and silvestrol vary between cell types. (A, B) Production of infectious virus progeny (PR8 strain) at 24 hpi was measured using plaque assay. The indicated cell lines were infected with MOI = 0.1 and treated with the increasing concentrations of pateamine A (A) or silvestrol (B) at 1 hpi. (C, D) Cell viability was measured using Alamar Blue assay after 24-h treatment with increasing concentrations of pateamine A (C) or silvestrol (D). Relative fluorescence values are normalized to vehicle control (DMSO). (A-D) Error bars represent standard deviations from 3 independent biological replicates.

**Figure 5.**
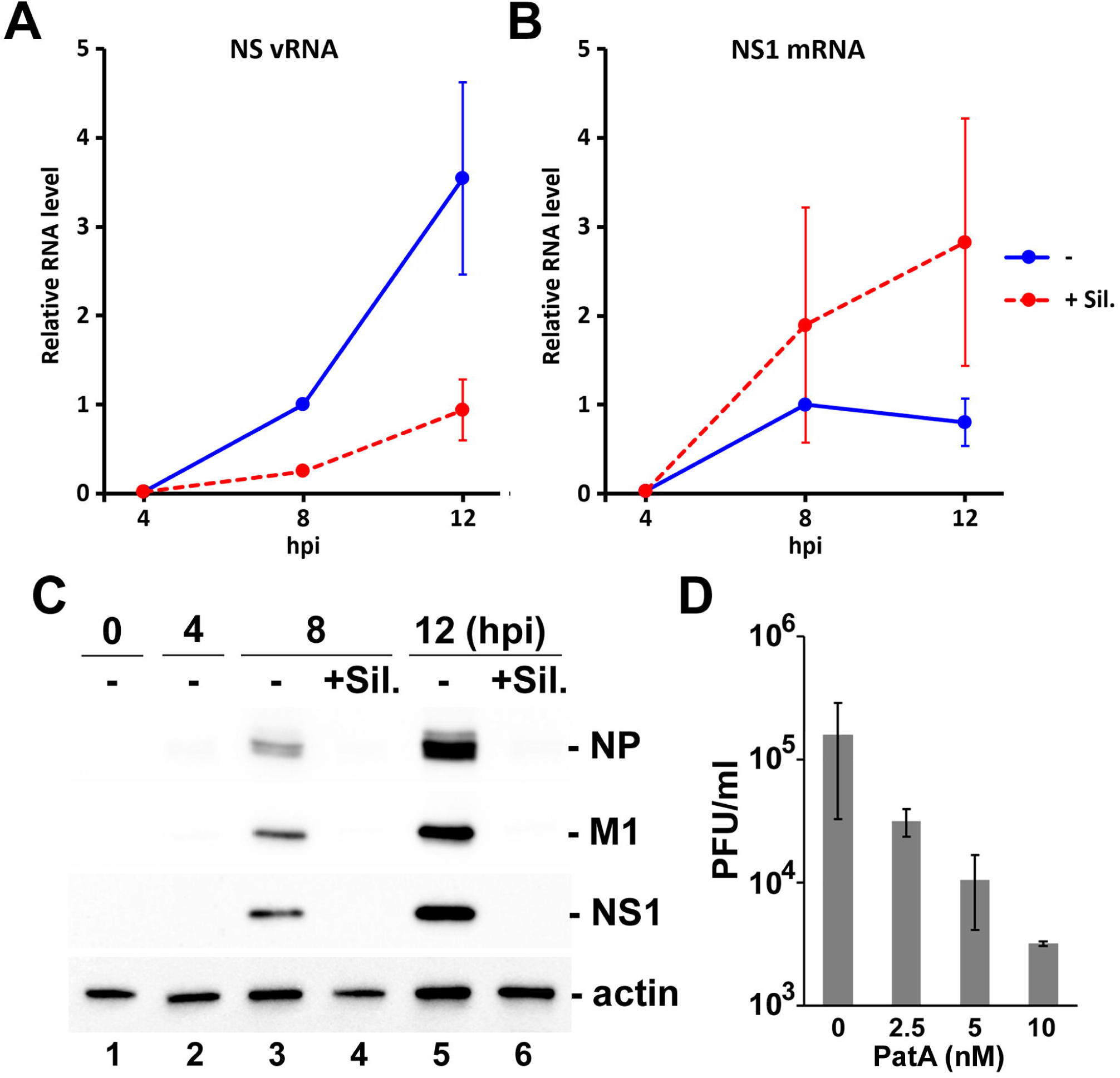
Silvestrol and pateamine A block replication of H3N2 strain of IAV. (A-C) A549 cells were infected with A/Udorn/72(H3N2) strain of IAV and treated with 320 nM silvestrol at 4 hpi. Total RNA and whole cell protein lysates were collected at 4, 8, and 12 hpi. The accumulation of viral NS segment vRNA (A) and NS1 mRNA (B) was measured using RT-qPCR, and the accumulation of viral proteins was analysed by western blotting (C). (D) Production of infectious virus progeny (Udorn strain) at 24 hpi was measured using plaque assay. A549 cells were infected with MOI = 0.1 and treated with the increasing concentrations of pateamine A at 1 hpi. Error bars represent standard deviations (n = 2).

### Silvestrol withdrawal causes SG dissolution and resumption of viral protein accumulation

PatA and silvestrol have distinct molecular structures and mechanisms of eIF4A disruption. Both molecules induce eIF4A dimerization and force an engagement with RNA, thereby depleting eIF4A from eIF4F complexes and inhibiting cap-dependent translation (14, 23). However, PatA has been shown to be an irreversible translation inhibitor (13), whereas silvestrol-dependent translation inhibition occurs reversibly (16). These distinct properties provided a unique opportunity to investigate whether IAV can ‘recover’ from SG formation following withdrawal of the SG-inducing drug. To address this directly, IAV-infected A549 cells were incubated with 30 nM PatA or 300 nM silvestrol starting from 1 hpi. At 4 hpi, drug was washed off of some infected cells, and SG formation and viral protein accumulation were analyzed using immunofluorescence microscopy at 12 hpi (Fig. 3A). Consistent with our previous observations (11), after withdrawal of PatA, SGs persisted for the remainder of the time-course, and viral proteins did not accumulate beyond the levels observed at 1 hpi. However, silvestrol withdrawal at 4 hpi led to SG disassembly and robust IAV protein accumulation; thus, ‘wash-off’ (WO) reversed the effects of silvestrol. To confirm that SG dissolution due to silvestrol withdrawal coincides with the resumption of protein synthesis, we treated A549 cells with silvestrol or PatA and labelled newly synthesized proteins with a 10 min pulse of puromycin at various times post-treatment (24). The newly-synthesized peptides were visualized by western blot (Fig. 3B). Both silvestrol and PatA caused strong inhibition of protein synthesis at 1 h post-treatment, which was sustained throughout the 24 h treatment time-couse. Silvestrol withdrawal at 3 h allowed for a complete restoration of protein synthesis at later times (Fig. 3B, lanes 7 and 12). By contrast, protein synthesis was never restored following PatA wash-off (Fig. 3B, lanes 8 and 13). Together, these findings demonstrate that IAV is able to recover from transient drug-induced translation arrest and resume the viral replication cycle when eIF4A inhibition is relieved.

**Figure 3.**
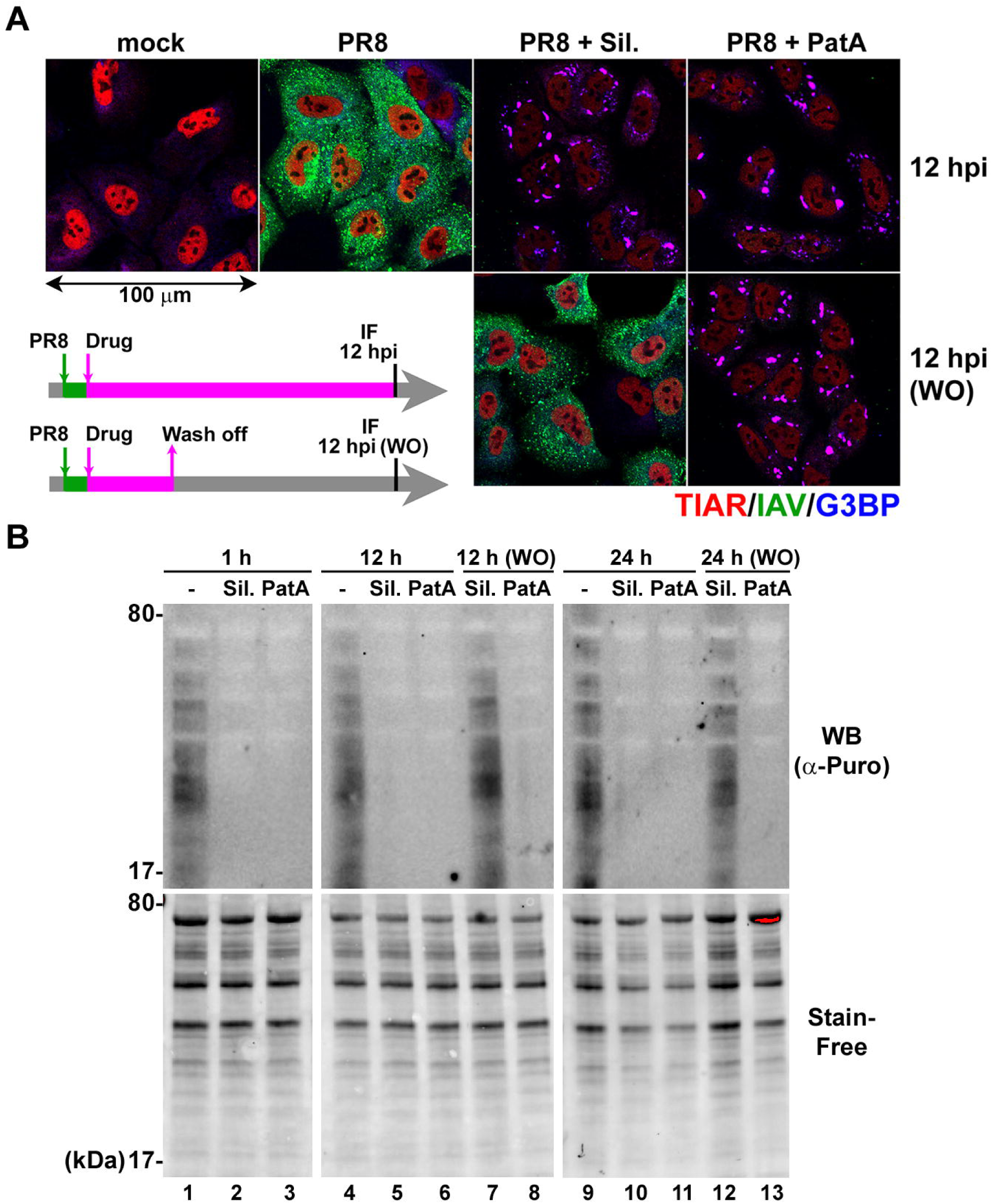
Translation inhibition by silvestrol is fully reversible. (A) A549 cells infected with PR8 strain of IAV were treated at 1 hpi with 320 nM silvestrol (Sil.) or 20 nM pateamine A (PatA) or left untreated. At 4 hpi, some wells were washed briefly with PBS and received fresh infection media without drugs as shown on the schematic outline of the experiment. At 12 hpi mock and virus-infected cells subjected to continuous incubation with Sil. or PatA or to drug wash off at 4 hpi (WO) were analysed by immunofluorescence staining with the polyclonal anti-influenza antibody (IAV, green) and the antibodies to SG markers TIAR (red) and G3BP (blue). (B) Total translation rates in A549 cells were analyzed using metabolic labelling with puromycin and subsequent western blotting with anti-puromycin antibody. In some cases, after initial 3-h treatments, Sil. or PatA were washed off (WO) prior to puromycin labeling. Total protein was visualized using BioRad Stain-Free reagent.

### Prolonged eIF4A inhibition triggers apoptosis

SG formation promotes cell survival in response to a variety of environmental stresses (9). However, the SG-inducing eIF4A inhibitors silvestrol and PatA have been shown to promote apoptosis of cancer cells (23, 25–27). Moreover, apoptosis is a well-described feature of late-stage IAV infection, required for efficient virion production (28–31). To determine whether eIF4A disruption affects the fate of IAV-infected cells in our system, both infected and uninfected A549 cells were treated with 300 nM silvestrol or 30 nM PatA, and apoptosis was measured by immunoblotting for PARP cleavage products. Following 16 h incubation with drugs, PARP cleavage species accumulated in both mock-infected and IAV-infected cells alike, indicating that sustained eIF4A disruption overcomes any pro-survival effects of SG formation, resulting in apoptosis (Fig. 4A, lanes 5-8). Interestingly, the sustained translation arrest by silvestrol not only diminished viral protein accumulation, but also prevented the RdRp from switching tasks between viral mRNA synthesis and genome replication (Fig. 4B). This finding is consistent with our previous observations for PatA (11), suggesting that both drugs affect the viral replication cycle in the same way. This dysregulated RdRp activity and an inability to produce the viral PKR antagonist NS1 due to translation arrest, enables detection of viral pathogen-associated molecular patterns (PAMPs), resulting in eIF2αphosphorylation (Fig. 4, lanes 5-6).

**Figure 4.**
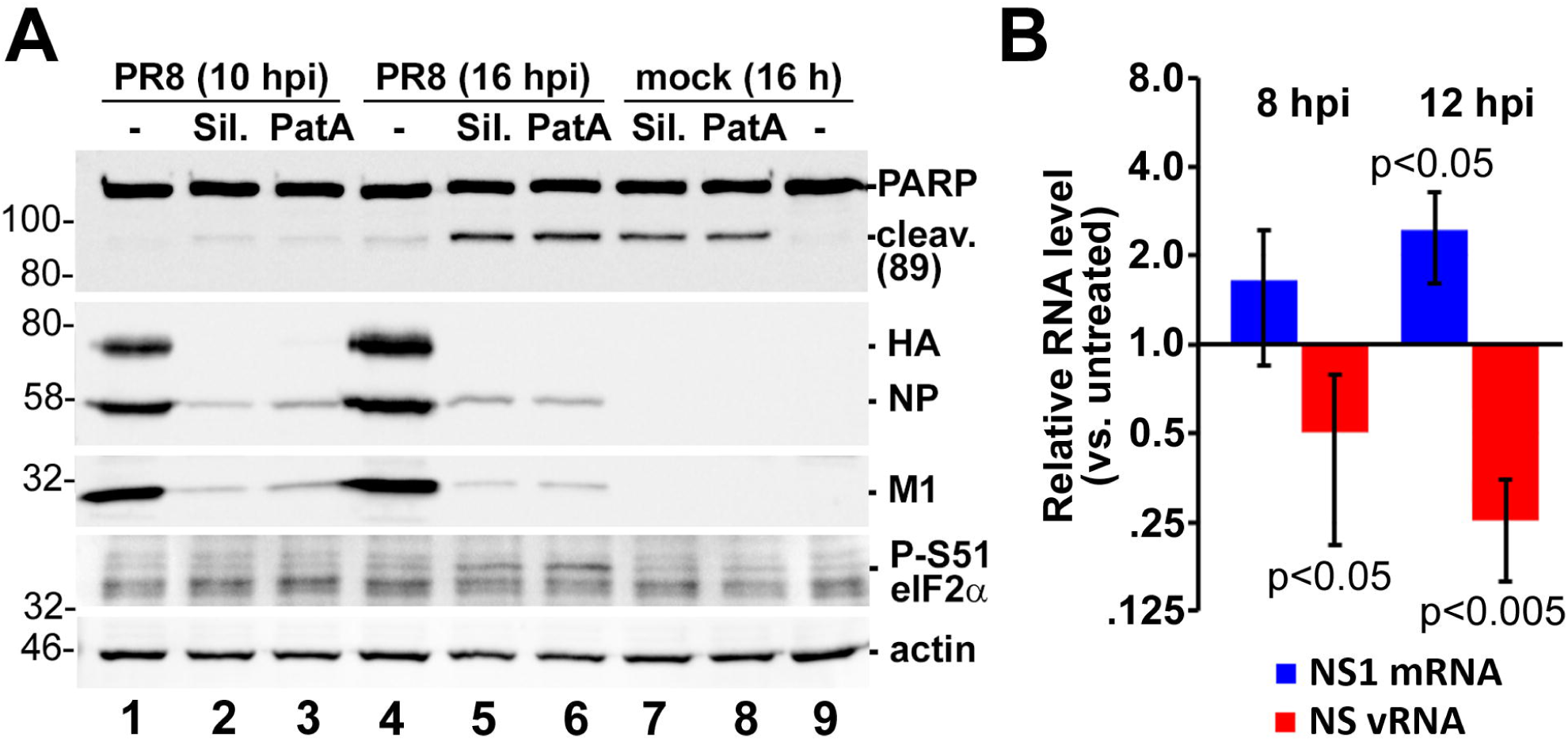
Sustained eIF4A inhibition by silvestrol and pateamine A leads to induction of apoptosis. (A) Western blotting analysis of A549 cell lysates obtained at the indicated times post-infection with the PR8 strain of IAV and treated with 320 nM silvestrol (Sil.) or 20 nM pateamine A (PatA) at 4 hpi or the equivalent time after mock infection. (B) Total RNA was isolated from cells treated with 320 nM silvestrol at 4 hpi and the relative levels of viral NS1 mRNA and vRNA at the indicated times post-infection were determined using RT-qPCR. Error bars represent standard deviations (n=3). P values were calculated using paired Student’s t-test.

### Effects of silvestrol and pateamine A on IAV replication are not strain-specific

We have determined that the lab-adapted H1N1 PR8 strain used in this study, which is the preferred backbone for vaccine production worldwide, is sensitive to eIF4A inhibition. To determine whether a genetically-divergent IAV strain could be similarly affected by eIF4A inhibiting drugs, A549 cells were infected with the H3N2 Udorn strain. Infected cells were treated with 300 nM silvestrol at 4 hpi, and the viral RNA and protein accumulation was determined over a 12 h time-course. Consistent with our previous findings using the PR8 strain, silvestrol treatment affected RdRp switching from viral mRNA synthesis to viral genome replication tasks (Fig. 5A, B) due to failure to accumulate viral proteins (Fig. 5C). Our findings to date indicate that PatA is better than silvestrol at inhibiting PR8 replication at sub-cytotoxic doses. For this reason, we investigated the effects of PatA on release of Udorn from A549 cells. We observed a similar magnitude of inhibition of Udorn virion production in cells treated with low nanomolar doses of PatA.

## DISCUSSION

IAV mRNAs generally resemble host mRNAs, which enables efficient translation by host cell machinery. However, these features also make them susceptible to stress-induced arrest of protein synthesis. Host translation initiation requires eIF4A, a helicase that unwinds mRNA secondary structure to permit ternary complex scanning for translation initiation codons. Here, we showed that IAV translation is sensitive to the eIF4A inhibitors PatA and silvestrol. These drugs limited viral protein accumulation and elicited the formation of SGs. Because progression through the viral replication cycle depends on accumulation of key viral proteins, these eIF4A inhibitors prevent the viral polymerase complex from switching from viral mRNA synthesis to viral genome replication. Both molecules could block replication of genetically-divergent IAV strains, PR8 and Udorn, suggesting a potential universal dependence on eIF4A activity. While the effects of silvestrol were reversible, PatA, known to bind irreversibly to eIF4A, sustained long-term arrest of viral protein synthesis following drug withdrawal.

Because many oncogenes have structured 5’-UTRs, and depend on eIF4A activity for their synthesis, eIF4A inhibitors like PatA and silvestrol, have been extensively studied for anti-cancer activity. Low doses of PatA have been shown to inhibit proliferation of tumor xenografts without appreciable toxicity in murine models (32). Indeed, PatA was able to inhibit oncogene synthesis at low doses that did not impinge on bulk protein synthesis rates, demonstrating that mRNA structure plays a crucial role in determining susceptibility and dose-dependent effects of eIF4A inhibitors. As in cancer cells, efficient virus replication typically demands sustained high rates of protein synthesis, which may likewise be dependent on eIF4A helicase activity. For example, Ebola virus has been shown to be exquisitely sensitive to eIF4A inhibition by silvestrol (20). Ebola virus mRNAs have highly-structured 5’-UTRs (33–35) and require processive helicase activity of eIF4A. By contrast, IAV mRNA 5’-UTRs are relatively short, and comprised of divergent host-derived mRNA segments fused to conserved viral mRNA segments. The heterogeneous nature of these 5’-UTRs challenges RNA structure prediction algorithms, but the short, conserved regions do not display significant secondary structure that would necessitate high eIF4A activity. Consistent with this, IAV mRNA translation is inhibited by relatively high doses of silvestrol and PatA that would be predicted to deplete eIF4A from translation preinitiation complexes. Our results and previous studies indicate that eIF4A helicase activity is required for translation initiation on IAV mRNAs, but it appears that processive unwinding of long, structured 5’-UTRs is not required. Consistent with this model, IAV infection was shown to deplete the eIF4A processivity factor eIF4B (36). The virus replicates efficiently in eIF4B-depleted cells, and likely benefits from diminished synthesis of eIF4B-dependent interferon-stimulated genes like IFITM3. Viral mRNP complexes likely lack eIF4B, but remain only partially characterized. They have properties that distinguish them from host mRNPs. For example, there is some evidence that eIF4E1 is also dispensable for viral mRNA translation (37). Moreover, IAV NS1 is known to stimulate viral mRNA translation, which may be linked to its ability to bind to viral mRNP complexes through interactions with eIF4G1 and PABP (38). A better understanding of the precise composition of viral mRNP complexes will likely inform our understanding of the role played by eIF4A and other core translation factors. Beyond these considerations of mRNP composition, dependence on eIF4A helicase activity might also be influenced by host shutoff, which is expected to markedly influence the availability of host translation factors.

We previously established that SGs do not form at any point during IAV infection (10), and that three viral proteins can inhibit SG formation (11). We also demonstrated that an early window of opportunity exists, before sufficient quantities of SG antagonizing viral proteins accumulate, when the virus is exquisitely sensitive to stress-induced translation arrest. Here, we further elucidated the mechanism of action of SG-inducing eIF4A inhibitors PatA and silvestrol. To date, most of our observations of SGs formed in IAV-infected cells indicate that these granules have canonical composition and properties. Throughout our studies, SG formation in infected cells reliably indicated disruption of viral protein accumulation and viral replication (10, 11). In this study, we observed that, upon withdrawal of silvestrol, SGs rapidly dissolved when viral protein synthesis resumed. At the same time, ongoing translation arrest triggered by PatA resulted in persistence of SGs throughout the observation period. It has been recently demonstrated that PKR is recruited to SGs by direct interaction with G3BP, and that this recruitment results in PKR activation (39, 40). We observe eIF2αphosphorylation in infected cells after prolonged SG induction by PatA or silvestrol. It will be interesting to determine whether this phosphorylation is dependent on PKR activation.

Our results show the magnitude of the threat that host-targeted translation inhibitors pose for viral replication. Nevertheless, the eIF4A inhibitors studied here have some undesirable properties that may be difficult to surmount. We observed that eIF4A inhibition resulted in cytotoxic effects that largely tracked with viral inhibition in all cell lines examined. The human (A549), dog (MDCK) and green monkey (Vero) cell lines studied here displayed markedly different sensitivities to silvestrol treatment, which cannot be explained by structural differences in eIF4A because all three isoforms of eIF4A (eIF4A-I, -II, and –III) are highly conserved between these species. Interestingly, while silvestrol has been shown to bind to eIF4A-I and eIF4A-II, PatA has been shown to bind to all three isoforms, including eIF4A-III (32, 41–43). While eIF4A-I and eIF4A-II function in the cytoplasm, eIF4A-III is localized to the nucleus and thus has no role in translation initiation. Instead, it is a member of the exon junction complex deposited on mRNA post-splicing, where it has been shown to participate in nonsense mediated decay (44). Moreover, eIF4AIII was previously shown to interact with the IAV polymerase complex (45). The functional significance of this interaction is unknown, and the contribution of eIF4A-III inhibition by PatA to its antiviral effects remain to be determined.

There is an outstanding need for new antivirals for influenza. Past history has shown that direct-acting antivirals are often rendered ineffective by rapid viral evolution. For this reason, host-targeted antivirals are an attractive alternative approach that should limit the emergence of drug-resistant variants. RNA silencing screens have shown that influenza virus replication depends on thousands of host genes, some of which may be potential candidates for therapeutic intervention. Our data indicates that inhibition of viral protein synthesis potently disrupts the viral replication cycle, and drugs that can block viral protein synthesis may serve as attractive candidates for host-directed antivirals. Despite their antiviral activity against IAV at high doses, PatA and silvestrol appear to lack specificity for viral translation complexes, and impede bulk translation in infected and uninfected cells alike. This distinguishes IAV from other viruses (e.g. Ebola, HCMV) and from cancer models, which were shown to be exquisitely sensitive to much lower doses of eIF4A inhibitor. For these reasons, an effective host-targeted antiviral translation inhibitor for influenza will ideally be specific for infected cells, while sparing uninfected cells. Alternatively, a detailed characterization of IAV mRNP complexes could highlight unique features that could be exploited by future antivirals.

## ACKNOWLEDGEMENTS

We thank members of the McCormick lab for critical reading of the manuscript. We thank Drs. Kevin Coombs (University of Manitoba, Winnipeg, MB, Canada), Yoshihiro Kawaoka (University of Wisconsin-Madison, Madison, WI, USA), Jerry Pelletier (McGill University, Montreal, QC, Canada), and Richard Webby (St. Jude Children’s Hospital, Memphis, TN, USA) for reagents. We thank Dr. Stephen Whitefield at the Dalhousie University Faculty of Medicine Cellular & Molecular Digital Imaging Core Facility for microscopy support. This work was supported by CIHR Operating Grants MOP-136817 and PJT 148727, and NSERC Discovery Grant RGPIN/341940-2012.

